# Green tea infusion aggravates nutritional status of the juvenile untreated STZ-induced type 1 diabetic rat

**DOI:** 10.1101/2020.01.13.904896

**Authors:** Luiz Carlos Maia Ladeira, Eliziária Cardoso dos Santos, Bruno Ferreira Mendes, Eliana Alviarez Gutierrez, Cynthia Fernandes Ferreira Santos, Fernanda Batista de Souza, Mariana Machado-Neves, Izabel Regina dos Santos Costa Maldonado

## Abstract

We have described for the first time the potential harmful effects of green tea on the metabolism and body composition of untreated juvenile experimental type 1 diabetic rats. The treatment containing 19.38% of epigallocatechin-3-gallate, its main catechin, increased blood glucose and water intake. It also increased oxygen consumption, enhanced energy expenditure and led to a lipid oxidation tendency in diabetic animals, which worsened the development of body fat in a way significantly more aggravated than diabetes alone. Taken together, our findings indicate that green tea treatment, when provided to juvenile diabetics, increases glycaemia, changes the body composition by reducing fat content and increases oxygen consumption, besides affecting energy expenditure. Therefore, the nutritional status of the juvenile type 1 diabetic rat is aggravated.

## 1. Introduction

Diabetes mellitus (DM) is a serious heterogeneous metabolic disease with increasing rates of incidence and prevalence worldwide ^1,2^. It is estimated that about 415 million adults have DM, and in 2040 figures are expected to reach 642 million ^3^. The disease-related complications, especially in type 1 DM (i.e. insufficient production of insulin by the pancreatic beta cells) include micro and macrovasculars disturbances ^4^, hepatic damage ^5^, renal ^6^, cardiac ^7^, and neurological impairment ^8^, aside from the poor nutritional status characterized by diabetes. Taken together, these complications become an important cause of morbidity and mortality, with negative impact on life quality, consequently reducing the life expectancy of these individuals ^1,3^.

Insulin therapy is effective and safe for the treatment of type 1 DM ^9^, but it alone does not eliminate the risk of complications from the disease. Therefore, non-pharmacological strategies, such as physical exercises and use of natural compounds with antioxidant polyphenols have been described as a complementary treatment ^10–14^. Within this category of compounds, studies have focused on the effect of epigallocatechin-3-gallate (EGCG), a catechin present in large amounts in green tea ^14–16^. Due to its potential therapeutic effect and pharmacological action (e.g. antioxidant, antidiabetic, anti-inflammatory and anti-apoptotic properties) described in previous studies ^15,17–20^, EGCG has been considered as a possible adjuvant in the treatment of diabetes, so as to improve the general health of individuals and, consequently, delay the development of DM complications ^13,14,21^. It has been suggested that this adjuvant profile is closely correlated with the inhibition of glucose production in hepatocytes ^22^. In fact, studies showed that green tea catequins suppressed hepatic gluconeogenic activity and activated the 5’-AMP-activated protein kinase (AMPK), which improved insulin signaling pathway and downregulated the genes that encode gluconeogenic enzymes ^22–24^.

Due to the antidiabetic activity described for EGCG and its antioxidant proprieties, various studies have explored the use of tea catechins, isolated or in combination with other drinks with similar proprieties, as an alternative to evaluate its systemic effect on different pathophysiological processes ^15,25^. Despite its relevant effects in most cases, one should consider the thermogenic potential of this kind of substance ^26,27^. It is well known that green tea polyphenolic compounds also modulate energy metabolism, thus enhancing thermogenesis, fat oxidation and energy expenditure ^27–30^. Therefore, caution should be taken, especially regarding the use of these substances to treat some diseases, since this type of response can be potentially noxious and aggravate an already installed pathological process.

In type 1 DM, some of the green tea metabolic effects can be harmful to a certain extent ^31–33^. Studies with experimental diabetes in rats showed that untreated diabetic animals present an impaired nutritional status and, especially when the disease appears in preadolescent rats or at younger ages, this condition can be aggravated and irreversible ^34,35^. Thus, the present study aimed to investigate the effects of green tea infusion on the nutritional status of type 1 diabetic rats, considering their feeding and murinometric parameters, body composition and metabolism of their untreated experimental model of type 1 diabetic rat.

## 2. Materials and methods

### 2.1. Animals and ethics statement

Eighteen male Wistar rats (30 days old; 82.52 ± 10.83g) were provided by the Central Animal Laboratory of the Center of Biosciences and Health from the Federal University of Viçosa. The animals were housed in polypropylene cages, in pairs, under controlled conditions of temperature (22 ± 2 °C) and light-dark cycles (12/12h). All animals received food (Presence Alimentos, Paulínea, SP, Brazil) and water *ad libitum*. The use of animals in the research was approved by the Ethics Committee of Animal Use of the Federal University of Viçosa (CEUA/UFV – protocol number 53/2018).

### 2.2. Preparation of green tea infusion

Five different lots of green tea (*Camellia sinensis*) leaves were obtained from Leão^®^ - Food and Beverages (Coca-Cola Company^®^). The lots were mixed (1:1) and the infusion was prepared mixing the leaves with warm distilled water (1:40 w/v, 80 °C) ^36^. The mixture remained infused for 20 minutes on a magnetic stirrer. Then, it was filtered through a 0.45 μm porous filter, frozen at −80 °C and lyophilized. The lyophilized samples were resuspended in distilled water at the moment of use.

### 2.3. Determination of total phenolic content

Total phenolic content was determined in triplicates as described before by Singleton and Rossi ^37^ using the Folin-Ciocalteau reagent. To that porpoise, an aliquot of 0.6 mL of the lyophilized extract resuspended in distilled water (1:25 w/v) was added to 3 mL of the Folin-Ciocalteau reagent. After 6 minutes, 2.4 mL of 7.5% sodium carbonate solution was added and agitated. The tubes were allowed to stand in dark for 1 hour at room temperature. The absorbance was measured at 760nm using an ultraviolet (UV)-spectrophotometer (BEL UV-M51, BEL Photonics, Italy). Different concentrations of gallic acid dissolved in distilled water were used to prepare the calibration curve (r^2^ = 0.9992). The total phenolic content was expressed as milligrams of gallic acid equivalent per gram of lyophilized samples of tea (mg GAE/g GTI).

### 2.4. EGCG analysis

EGCG analysis was performed as described by Kim-Park et al.^38^, with some modifications. High-performance liquid chromatography (HPLC) (Prominence LC-20A, Shimadzu, Kyoto, Japan), equipped with Diode Arrangement Detector (DAD), LC-20AD pump, SPD-M20A detector, CTO-20A oven and LabSolutions software, was used to determine the EGCG content using a maximal absorption peaks at 272nm. It was used a Vydac C18 (4.6 × 250 mm) column, at 30 °C, with a 5μL injection volume. The mobile phase was composed of water and 2.0% acetic acid (1:1). The infusion lyophilized powder was suspended in methanol before analysis. The mobile phase flow rate was 1.0 mL/min and the run time was 15 min. The retention time of EGCG was 4.5 min and the total amount of it was calculated using a standard curve (r^2^ = 0.9967) developed under the same conditions using an EGCG chemical standard (≥ 98.0%, Sigma Aldrich Inc. - CAS Number 989-51-5. St. Louis, MO, USA).

### 2.4. Antioxidant capacity by the 2,2’-Azinobis-[3-ethylbenzthiazoline-6-sulfonic acid] (ABTS) decolorization assay

ABTS radical stabilization by the lyophilized tea extract was determined at a wavelength of 734 nm on an ultraviolet (UV)-spectrophotometer (BEL UV-M51, BEL Photonics, Italy), following the described method by RE et al.^39^. Different concentrations of trolox dissolved in ethanol (80 %) were used to prepare the calibration curve (r^2^ = 0.9996). The antioxidant capacity by the ABTS method was expressed as μMol of trolox equivalent per gram of lyophilized samples of tea (μMol TE/g GTI).

### 2.5. Antioxidant capacity by ferric reducing antioxidant power (FRAP) assay

The FRAP assay was performed as described before ^40^. An aliquot of the lyophilized extract resuspended in distilled water (1:25 w/v) was added to the FRAP reagent and incubated at 37°C for 30 min. The absorbance was determined at a 595 nm wavelength on an ultraviolet (UV)-spectrophotometer (BEL UV-M51, BEL Photonics, Italy). Different concentrations of ferrous sulphate (FeSO_4_) dissolved in distilled water were used to prepare the calibration curve (r^2^ = 0.9985). The antioxidant capacity by the FRAP method was expressed as μMol of FeSO_4_ equivalent per gram of lyophilized samples of tea (μMol FeSO_4_/g GTI).

### 2.6. Experimental design

After seven days of acclimation in the bioterium, six rats were randomly selected to integrate the healthy control group. Type 1 diabetes was induced in 12 rats, after 12 h fasting by a single intraperitoneal (i.p.) injection of streptozotocin (STZ) (Sigma Chemical Co., St, Louis, MO, USA) at a dosage of 60mg/Kg of body weight (BW) diluted in 0.01 M sodium citrate buffer, pH 4.5. The control group received the buffer alone by the same administration route ^12^. Two days after the STZ injection, following 12 h fasting, blood samples were collected from the tail vein and glycaemia was measured using a glucometer (Accu-Chek® Performa, Roche LTDA). All animals presented fasting blood glucose levels higher than 250 mg/dL and were included in the study. The hyperglycemic rats were divided into two groups (n = 6, each). Thus, the experimental protocol consisted in three groups: healthy control group (Ctrl, n = 6); diabetic control group (Diabetes, n = 6); and the diabetic group treated with the green tea infusion (GTI diabetic, n = 6), which received a 100mg/Kg dosage of GTI, diluted in 0.6mL of water. The control groups received 0.6mL of water alone. All treatments (GTI and water) were administered by gavage, every day, during 42 days.

Considering that type 1 diabetes usually appears at young ages ^41^, the experimental protocol started when the animals were 40 days old and finished when they reached 82 days of age, (i.e. from periadolescence to the early adult phase) ^42^.

After the experimental protocol period, the animals were euthanized by deep anesthesia (sodium thiopental, 60mg/Kg i.p.) followed by cardiac puncture and exsanguination ^33^.

### 2.7. Blood glucose, body weight and food and water consumption

Fasting blood glucose was measured in blood samples from the tail vein using a glucometer and reactive strips (Accu-Chek® Performa, Roche LTDA. Jaguaré, SP, Brazil). Body weight, water and food consumption were measured using a precision scale (BEL M503, 0.001g, Piracicaba, SP, Brazil). All these parameters were monitored weekly.

### 2.8. Murinometric and feeding parameters

All the murinometric and feeding measurements were calculated as described by Nery et al.^43^. On the last day of the experimental protocol, the naso-anal length (NAL) of the rats was measured with an inelastic measuring tape (e = 0.1 cm) to calculate the following indicators: Lee index 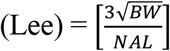and Body Mass Index 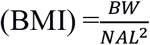, where BW refers to the final body weight, and NAL, the naso-anal length. The following feeding indexes were also calculated: Specific Rate of Weight Gain 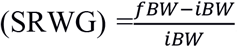, where iBW refers to body weight at the beginning of the experiment and fBW refers to the final BW. Feeding efficiency was indicated by the Coefficient of Feeding Efficiency (CFE) and Weight Gain per Caloric Consumption (WGCC), calculated as follows: 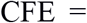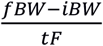, where tF refers to the total amount of food ingested (g) in the experiment. 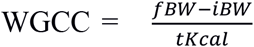 where tKcal stands for the total amount of Kcal ingested in the experiment.

### 2.9. Dual-energy X-ray absorptiometry analysis

Body composition was evaluated under anesthesia (sodium thiopental, 60mg/Kg i.p.) on the 36^th^ day of treatment. The rats were positioned in ventral recumbency on the scan table. All scans were performed using dual-energy X-ray absorptiometry (DXA) (Lunar, DPX, Madison, WI, USA) to evaluate fat (% and g) and lean mass (%). An accelerating voltage of 100 kV with current of 0.188 mA and radiation dose of 10 microGy were used for scanning. The Encore v.13 2011 (GE Healthcare Systems, Chicago, IL, USA) software system was used for data analysis. The results were expressed as a mean value.

### 2.10. Calorimetric analysis

The oxygen (O_2_) consumption and carbon dioxide (CO_2_) production of the experimental animals were measured through gas analyzer (Oxyleptro, Harvard Apparatus, Holliston, MA, USA) on the 40^th^ day of treatment, without fasting. To that end, the ambient air was pumped through a metabolic chamber and samples of the extracted air were directed to the gas analyzer (air flow = 1.0 L/min). The Metabolism (Panlab, Barcelona, Spain) software system was used for data analysis. The animals remained for 60 minutes in the metabolic cage mimicking their real conditions in the laboratory for the determination VO_2_ (mL/min/Kg^0.75^) and VCO_2_ (mL/min/Kg^0.75^) at rest. The test was performed with animals from the three experimental groups, concomitantly, from 6 pm to 11 pm ^44^.

The respiratory quotient (RQ) and the total 24 h energy expenditure rate (EE) (Kcal/day) were calculated using the following equations: RQ = VCO_2_/VO_2_, where VCO_2_ refers to the volume of CO_2_ produced by the rats and VO_2_, the O_2_ volume consumed during the assay; and *EE* = (3.815 + (1.232 ∗ *RQ*)) ∗ *VO*_2_ ∗ 1.44 is used for energy expenditure.

### 2.11. Statistical analysis

All the results were submitted to the Shapiro-Wilk test for normality assessment. The data expressed as percentage were transformed by angular transformation before the analysis. The results were expressed as mean ± standard deviation (mean ± SD) and analyzed using unpaired Student’s *t*-test when the variances were equal (by *F* test) and unpaired Student’s *t*-test with Welch’s correction for data with unequal variances (Ctrl vs Diabetes; Diabetes vs GTI diabetic). Statistical significance was established at *P* ≤ 0.05. All tests and graphics were performed using the GraphPad Prism 6.0 statistical software system (GraphPad Software Inc., San Diego, CA, USA).

## 3. Results

### 3.1. Green tea infusion phytochemical analysis

The total amount phenolic components in the green tea infusion lyophilized powder was evidenced to be 3.88 ± 2.49 mg GAE/g GTI. The EGCG content, analyzed by HPLC methodology, was shown to be 19.38% of the total GTI content. The extract presented an antioxidant capacity of 3.26 ± 0.06 μMol TE/g GTI in the ABTS assay and 46.38 ± 4.1 μMol FeSO_4_/g GTI in the FRAP assay.

### 3.2. Green tea infusion increases glycaemia and favors the polydipsia in diabetic animals

After diabetes induction and subsequent hyperglycaemia confirmation in the experimental animals (i.e. above 250 mg/dL), both diabetic groups maintained high blood glucose levels, which remained above 400 mg/dL, compared with the healthy control group (glucose < 100 mg/dL) in the last four weeks of the experimental protocol. Besides, GTI diabetic rats presented glycemic levels significantly aggravated (*P* = 0.0223; Fig. 1 B). Increased glucose levels were consistent with the consequent increment in water consumption in the same experimental groups (Fig. 1 D). The diabetic animals maintained high water consumption compared to the Ctrl group during the entire experiment. In the sixth week, this value was significantly higher for the GTI diabetic group (*P* = 0.0296), compared with the diabetic animals that did not receive the green tea infusion and between the diabetic group compared with the healthy control group (*P* < 0.0001; Fig. 1 D).

**Fig. 1.**
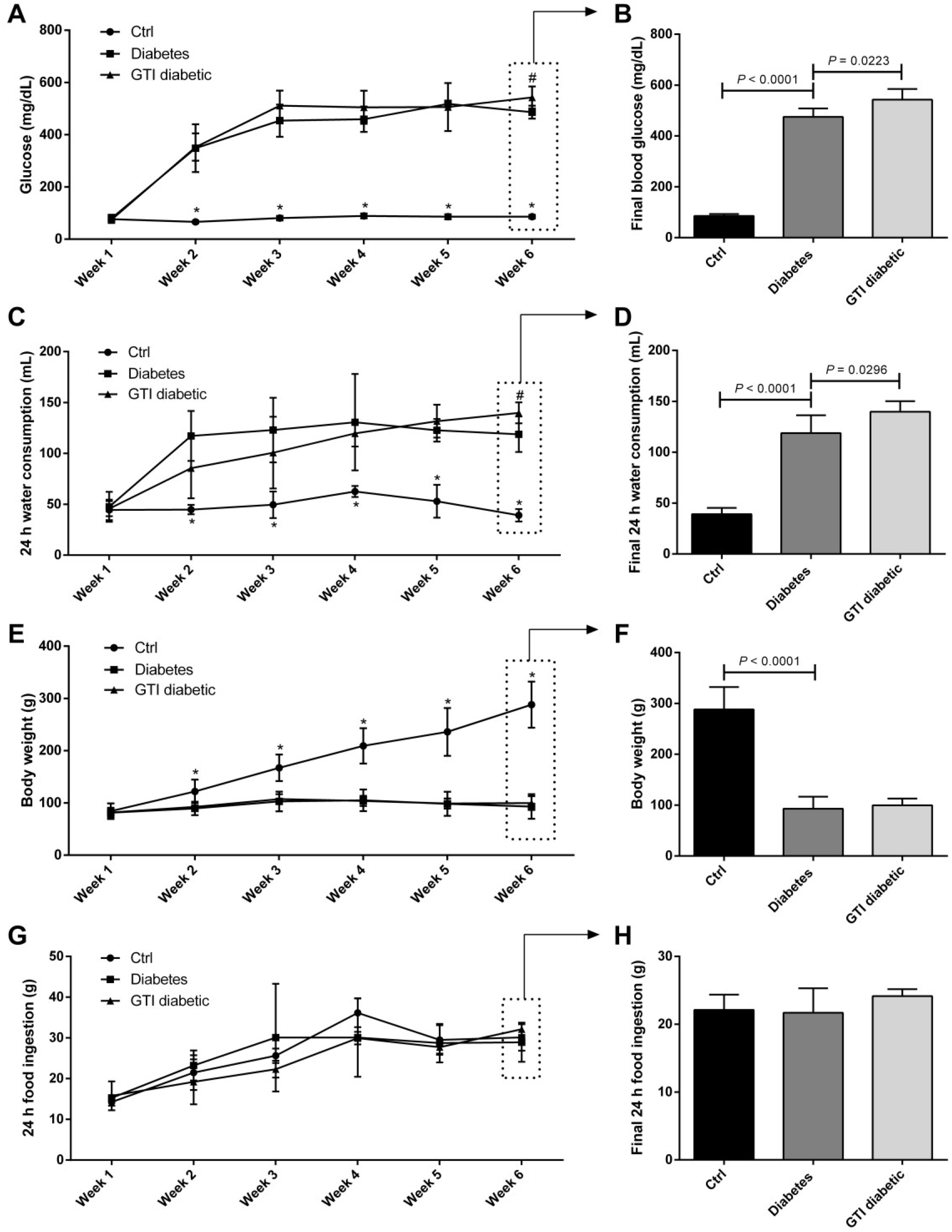
Body weight, food and water intake, and glucose levels of male Wistar healthy and diabetic rats treated with green tea infusion. **A -** 12h fasting blood glucose (mg/dL). **C -** Daily average water consumption (mL). **E -** Body weight (g) measured weekly. **G -** Daily average food ingestion (g). The **B**, **D**, **F** and **H** graphs represent the same variables featured in the last week of the experiment. Mean ± SD. In the A, C, E and G graphs, the asterisk (*) indicates that Diabetes group is statistically different from Ctrl, and the hash (#) indicates that GTI diabetic is different from the Diabetes group. The statistical differences are indicated with bars at the B, D, F and H graphs, with the *P* value above the bars. The data were compared (Ctrl vs Diabetes; Diabetes vs GTI diabetic), considering statistical differences when *P* ≤ 0.05. (n = 6 animals/group).

Body weight (Fig. 1 E) presented normal evolution during the six weeks of the experimental protocol for the healthy control group, ranging from 100 to 280g from the first to the sixth week, respectively. In the diabetic animals, this weight gain was severely impaired and no additional weight was registered in these animals during the six weeks of the experiment. This deficiency was observed in the sixth week (Fig. 1 F). Body weight gain decreased significantly in diabetic animals (*P* < 0.0001), compared to the control. No differences for this variable were observed between the diabetic groups, treated with GTI or not. No statistic differences were observed for food ingestion between the control and the experimental groups (Fig. 1 G and H) during the six weeks of the study.

### 3.3. Diabetes changes murinometric and feeding parameters

The murinometric parameters indicate that the rats from the diabetic groups remained with similar body proportions, but were different from the healthy control group, as indicated by NAL and the BMI (*P* < 0.0001; Table 1). The Lee index shows that body weight is proportional to NAL in all groups, regardless of the rat size.

**Table 1:** Murinometric and feeding parameters of male Wistar healthy and diabetic rats treated with green tea infusion

|  | Ctrl | Diabetes | GTI diabetic |
| --- | --- | --- | --- |
| Naso-anal lenght (cm) | 22.29 ± 1.25 | 15.60 ± 1.34* | 16.00 ± 1.27 |
| Lee index (g/cm) | 0.29 ± 0.01 | 0.29 ± 0.01 | 0.29 ± 0.03 |
| BMI (g/cm<sup>2</sup>) | 0.55 ± 0.06 | 0.40 ± 0.06<sup>#</sup> | 0.40 ± 0.09 |
| SRWG (g/Kg) | 2.44 ± 0.25 | 0.15 ± 0.26* | 0.23 ± 0.20 |
| CFE (g/g food) | 0.185 ± 0.019 | 0.009 ± 0.018* | 0.017 ± 0.015 |
| WGCC (g/Kcal food) | 0.048 ± 0.005 | 0.002 ± 0.004* | 0.004 ± 0.004 |

The data (Mean ± SD) were compared (Ctrl vs Diabetes; Diabetes vs GTI diabetic) considering statistical differences when $P \leq 0.05$ . (n = 6 animals/group). Asterisk (\*) indicates difference between the Ctrl and Diabetes group ( $P < 0.0001$ ), and the hash (#) indicates different means from the same comparison ( $P = 0.0049$ ). BMI – body mass index; SRWG – specific rate of weight gain; CFE – coefficient of feeding efficiency; WGCC - weight gain per caloric consumption.

Food intake did not differ throughout the experimental period and between experimental groups. However, the differences in BW and murinometric parameters can be explained by the feeding efficiency parameters. SRWG, CFE, and consequently WGCC, presented lower values in the diabetic groups (*P* < 0.0001; Table 1F), which indicates decreased efficiency in the conversion of food nutrients into tissue components.

### 3.4. Green tea infusion reduces fat mass gain in diabetic animals

The body composition examination revealed significant differences in the fat mass of the diabetic animals in both percentage levels (%) and absolute amount (g) when compared with the control animals (*P* < 0.0001; Fig. 2 A and B, respectively). Green tea infusion accelerated this response by significantly reducing fat accumulation, which is demonstrated by the relative amount of fat at the end of the experiment (*P* = 0.0045).It reached 9.3 ± 2.9% of the total body weight with the minimum value of 5.8%. Similar behavior was observed in its absolute amount compared to the untreated diabetic animals (*P* = 0.0053). Consequently, the relative lean mass (Fig. 2 C) represented a major portion of the body of the rats in the Diabetic group (*P* < 0.0001) compared to the Ctrl, and even higher in the GTI diabetic (*P* = 0.005) compared to the Diabetes group.

**Fig. 2.**
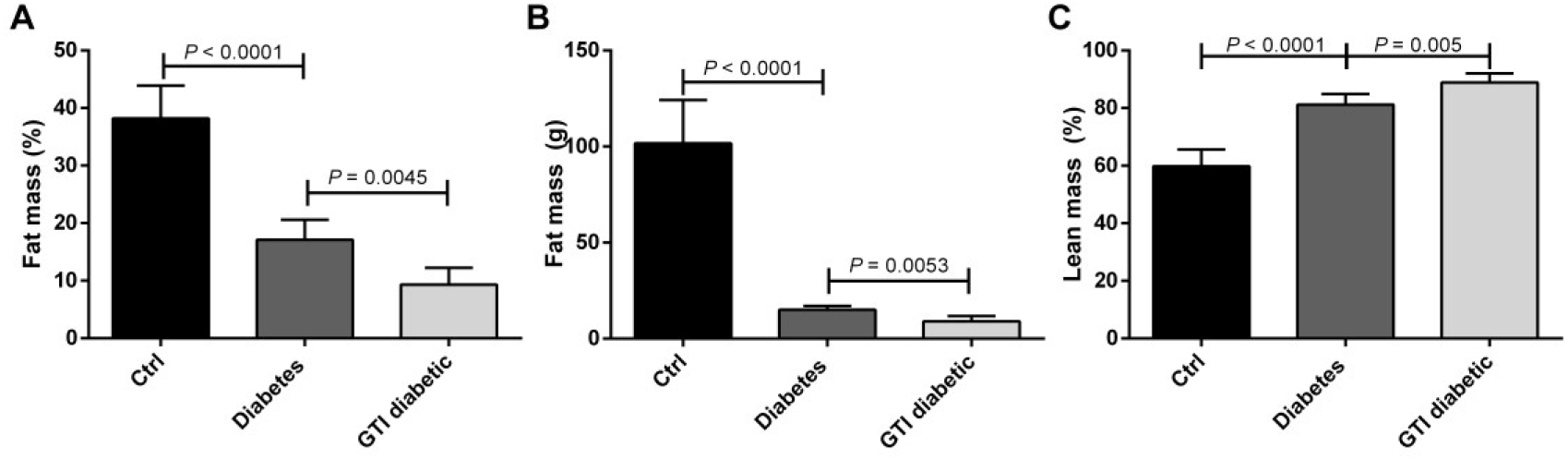
Body composition of male Wistar healthy and diabetic rats treated with green tea infusion. **A** – Relative fat mass (%). **B** – Absolute fat mass (g). **C** – Relative lean mass (%). The data are represented as Mean ± SD. The statistical differences are indicated with bars in the graphs, with the p value above the bars. The data were compared (Ctrl vs Diabetes; Diabetes vs GTI diabetic) considering statistical differences when *P* ≤ 0.05. (n = 6 animals/group).

### 3.5. Green tea infusion elevates the energy expenditure of diabetic animals

The calorimetric analysis revealed modulation in the metabolism of diabetic rats. Increased oxygen consumption was observed in the Diabetes group (*P* = 0.0058; Fig. 3 A) resulting in higher energy expenditure throughout the day (*P* = 0.0106; Fig. 3 D). On the other hand, animals treated with the GTI presented higher oxygen consumption and EE, when compared with the diabetic group that received the placebo treatment (water) (*P* < 0.05; Fig. 3 A and D). As shown in Fig. 3 C, the respiratory quotient (RQ) of the diabetic groups was lower, compared to the results of the control group (*P* = 0.0010), which indicates variation in the preferential macronutrient substrate used for energy generation.

**Fig. 3.**
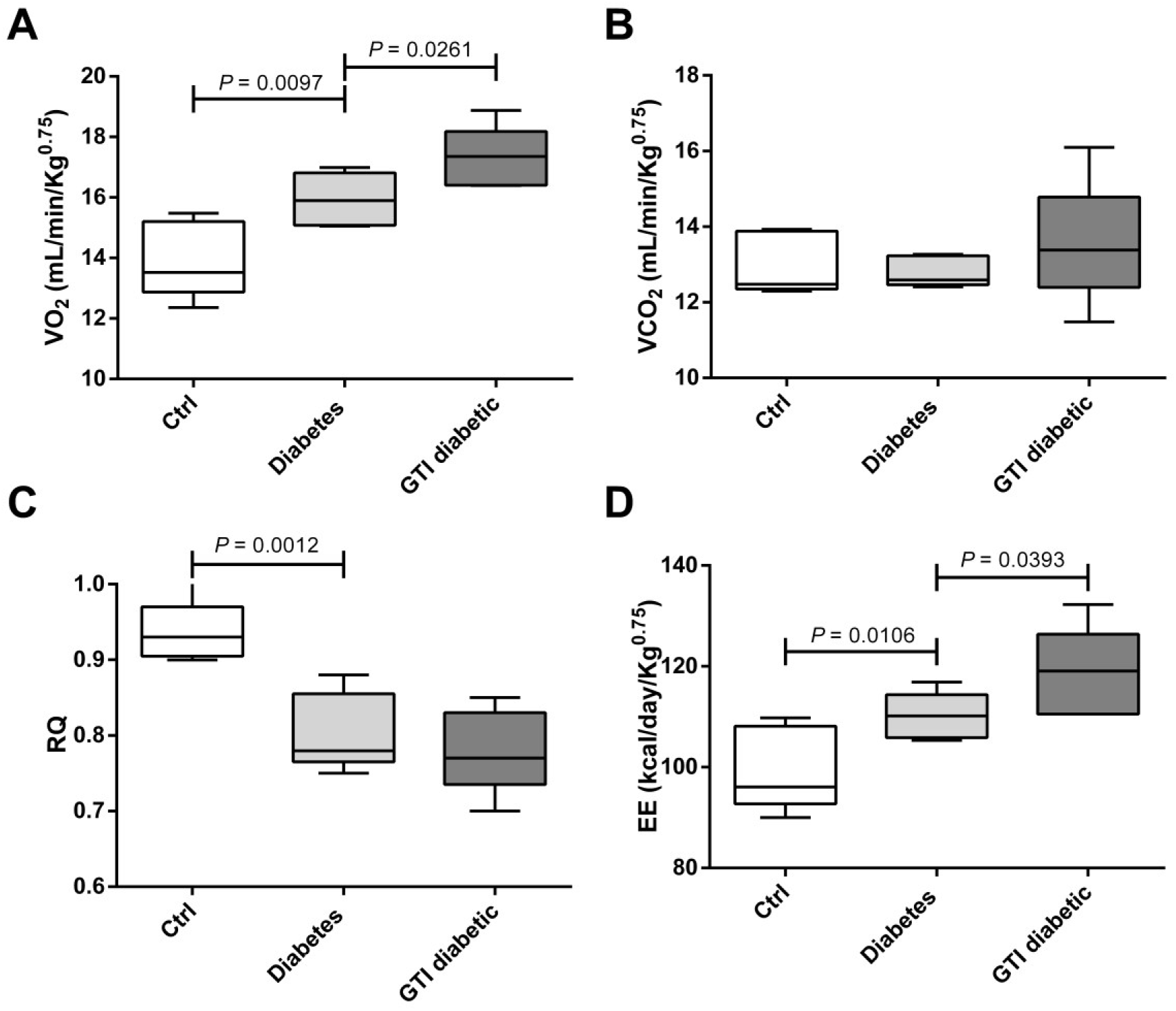
Calorimetric analysis of male Wistar healthy and diabetic rats treated with green tea infusion. **A** - VO_2_, average volume of oxygen consumed (mL/min/Kg^0.75^). **B** - VCO_2_, average volume of carbon dioxide produced (mL/min/Kg^0.75^). **C** - Respiratory quotient. **D -** Daily energy expenditure (EE) (Kcal/day/Kg^0.75^). The data are presented as Mean ± SD. The box represents the interquartile interval with the mean indicated (horizontal line), and the whiskers represent the superior and inferior quartiles. The statistical differences are indicated with bars in the graphs, with the *P* value above the bars. The data were compared (Ctrl vs Diabetes; Diabetes vs GTI diabetic) considering statistical differences when *P* ≤ 0.05. (n = 6 animals/group).

## 4. Discussion

Large amount of catechins are found in green tea (i.e. obtained from the *Camellia sinensis* L.). Their effects have been extensively explored ^45–47^ due to their potential health benefits in the treatment and prevention of human diseases. Its antidiabetic properties have been proven in previous studies ^17–19^, but this is the first time that strong evidence is reported that green tea infusion has great impact on glycaemia, body composition, nutritional status and metabolic activity in young streptozotocin-induced diabetic rats.

Admittedly, regarding the hypoglycemic effects of green tea and its components, specifically EGCG, important therapeutic potential has been demonstrated under experimental conditions ^1,14,15,48,49^. This, in turn, has been closely related to the potential of this substance to increase insulin activity ^19,50^. However, it was observed that green tea containing a proven amount of 19.38% of EGCG contributed differently in that parameter. The diabetic animals treated with green tea presented considerably higher blood glucose levels compared to the untreated rats. Streptozotocin, used in our study to induce the diabetic condition in the experimental animals, destroys the pancreatic insulin producer beta cells, leading to hyperglycemic condition ^51^. Thus, a positive relation cannot be attributed to the interaction between tea catechins and insulin. Therefore, it is possible to hypothesize that the maintenance of hyperglycaemia by GTI in our investigation may be related to the differential expression of glucose transporters or alterations in energetic metabolic pathways.

Aligned with this perspective, Kobayashi et al.^52^ showed that green tea catechins can inhibit the sodium-dependent glucose transporter 1 (SGLT1) in the brush-border membrane of enterocytes. This *in vitro* study, using brush-border membrane vesicles obtained from the small intestine of healthy rabbits, demonstrates that catechins containing the galloyl radical (epicatechin gallate and EGCG) were shown to bind to SGLT1, rendering the transporter unusable. Due to the inhibition of glucose uptake by the intestine and consequent drop in blood glucose, such an outcome could be encouraging. However, animals with STZ-induced diabetes are able to express another glucose transporter at the brush-border and basolateral membranes of enterocytes, such as transporter GLUT1, which is not expressed on the brush membrane of enterocytes in healthy animals ^53^, and GLUT2, that is inserted in the brush-border membrane when the luminal amount of glucose is still significant ^54–56^. GLUT1 expression, combined with GLUT2 regulation, maintains intestinal glucose uptake, regardless of the inactivation of SGLT1, thus preserving the hyperglycemic condition of the STZ-diabetic animal.

It is well established that persistently elevated glycaemia exacerbates the symptoms of type 1 diabetes (i.e. polyuria, polyphagia, polydipsia, extreme fatigue, weight loss despite high food intake), as reviewed by Ullah et al.^8^. Green tea contributed to aggravate these symptoms, since tea in our study reinforces the maintenance of hyperglycaemia. Although polyphagia is also a symptomatic consequence of type 1 diabetes ^8^, all groups presented similar food consumption throughout the experiment. On the other hand, body weight was compromised in diabetic animals, regardless of GTI intake. In contrast, studies were consistent in showing a positive relationship between green tea consumption and increased body weight gain in untreated experimental type 1 diabetes ^13–15^. However, in these studies, diabetes was induced in adult animals already presenting an optimal level of body development. We believe that the fact that diabetes was induced in animals at periadolescent age has caused the poor development and compromised weight gain observed during the six weeks of study.

Those differences in body weight, combined with similar levels of food consumption generated lower values of feeding efficiency features in our experimental model. Thus, the specific rate of weight gain as well as the coefficient of feeding efficiency and weight gain per caloric consumption were significantly lower in diabetic groups regardless of green tea consumption. Aligned with this perspective, other studies had already shown clear evidence that confirm these findings ^57–59^. These data reveal reduced efficiency in food nutrient conversion into tissue components, as previously described ^35^. Herrero et al.^60^ attributed this fact to the lack of plasma insulin, which prevents the transport of glucose to insulin-dependent cells (i.e. adipocytes, myocytes and cardiomyocytes), thereby forcing changes in metabolic routes so as to increase fat use.

This impairment in weight gain also delayed the body development of the diabetic rats. This finding can be corroborated by both the growth impairment of their naso-anal length and the stagnant fat mass accumulation observed in the X-ray absorptiometry scanning. According to Silva et al.^34^, when diabetes occurs at young age, it may compromise normal bone development. We did not directly evaluate this tissue, but some studies consistently point out the positive relation between diabetes and poor bone mineral metabolism and consequent impaired animal growth ^34,35^. This condition impacts the rate of bone mineral apposition and decreases the activity of osteoblastic cells, which leads to premature bone growth interruption, with consequent impairment to bone development in murine models of type 1 diabetes induced by STZ or alloxan. These facts consequently impair the length and size development of diabetic animals ^34,35^. We also proved their negative impact on the body composition of diabetic animals, in which impaired fat mass gain was aggravated when green tea was administered.

It has been discussed the relation between the EGCG, present in green tea, and increased lipolysis secondary to glucagon secretion ^61,62^. Studies have shown that EGCG is a potent inhibitor of the enzyme catechol-o-methyltransferase (COMT), which degrades norepinephrine ^63,64^. Norepinephrine persistence maintains beta adrenergic stimuli in pancreatic alpha cells, which increases glucagon production and release ^65^. Without inhibition by insulin, glucagon stimulates glycogenolysis in the liver until the depletion of the glycogen stocks ^61^. At this point, glucagon also stimulates gluconeogenesis, leading to the production of glucose from other substrates, such as proteins, besides increasing lipolysis and reducing fat deposits ^61,66^. Although we have not quantified glucagon, these mechanisms can explain the green tea impact on body composition in diabetic animals.

This persistent activation of the beta adrenergic stimuli mediated by green tea catechins in the pancreatic alpha cells, as previously described, corroborates the findings of higher oxygen consumption (VO_2_) and daily energy expenditure (EE: Kcal/day/Kg^0.75^) in diabetic animals treated with tea in our study. Type 1 diabetes induces higher oxygen consumption by modulating the energetic metabolism and the substrate utilization in energy production ^60,67^. Qualitatively, the respiratory quotient (RQ) indicates the types of energy substrate the animal preferentially consumes. Our control animals presented an RQ ranging between 0.9 and 1.0, which indicates a preference for carbohydrate hydrolysis. On the other hand, diabetic groups presented an RQ between 0.7 and 0.9, which indicates major fat oxidation for energy production ^68–70^. Tea catechins are linked to an improved expression of proteins related to beta oxidation and thermogenic capacity ^27,71,72^. Both mechanisms require an expanded mitochondrial activity that, in turn, leads to an increased demand of oxygen ^8,73^. We measured and demonstrated that the diabetic animals consumed more oxygen than the healthy control group. The green tea treatment, in contrast, increased oxygen consumption, which reflected in the daily energy expenditure of the rats treated with tea, maybe due to increased metabolic rate and/or the stimulation of lipolysis, beta oxidation and thermogenesis.

Treating young type 1 diabetic animals with green tea or its catechins seems to be a two-way pathway. At first, the use of tea and its molecules with highly antioxidant capacity seems effective against diabetes complications, as exhaustively described by the scientific literature. However, these molecules have other activities. They increase the mobilization and use of fat as energy source by the organism and even stimulate thermogenesis, processes that generate large amounts of reactive oxygen species. Catechins also contribute to maintain the hyperglycaemia by glycogenolysis and gluconeogenesis stimulated by glucagon. The most likely explanation for the lack of hypoglycemic effect of green tea combined with the impaired fat mass gain and increased energy expenditure in this study seems to be the hypothesis of COMT inhibition with consequent prolongation of beta adrenergic pathway stimulation in pancreatic alpha cells and thermogenic adipocytes.

In experimental type 1 diabetes not treated with insulin in young animals, the effect of green tea remains controversial. Even with the previously reported beneficial effects, these results are subject to factors such as the age at which the disease is induced. Collectively, we propose that 1) the studied parameters behave differently when observed in animals with type 1 diabetes induced at periadolescence or younger ages, when the disease is aggravated. 2) When diabetes appears at the juvenile ages, the green tea treatment increases glycaemia, changes body composition by reducing the fat content and increases oxygen consumption. It affects energy expenditure and worsens the nutritional status of the young type 1 diabetic rat.

## Acknowledgements

The authors are thankful to Farias, L. M. of the Laboratory of Biodiversity of the UFV for the chemical analysis in HPLC of the green tea extract; Dominik, D. J. S. for the assistance with logistics and the experiment; Magalhães, C. O. D. for the DXA scanning; Silva, M. R. for the statistical insights; Sacramento, E. for the English language review; da Silva, J. for the manuscript review and Coordenação de Aperfeiçoamento de Pessoal de Nível Superior (CAPES) for the L. C. M. Ladeira Ph.D. scholarship (Procs. Nr. 88882.436984/2019-01).

## References

1. Al Hroob AM, Abukhalil MH, Hussein OE, Mahmoud AM. Pathophysiological mechanisms of diabetic cardiomyopathy and the therapeutic potential of epigallocatechin-3-gallate. Biomed Pharmacother. 2019;109(October 2018):2155–2172. doi:10.1016/j.biopha.2018.11.086.

2. World Health Organization. Global Report on Diabetes. Geneva; 2016.

3. Ogurtsova K, da Rocha Fernandes JD, Huang Y, et al. IDF Diabetes Atlas: Global estimates for the prevalence of diabetes for 2015 and 2040. Diabetes Res Clin Pract. 2017;128:40–50. doi:10.1016/j.diabres.2017.03.024.

4. Chawla A, Chawla R, Jaggi S. Microvasular and macrovascular complications in diabetes mellitus: Distinct or continuum? Indian J Endocrinol Metab. 2016;20(4):546. doi:10.4103/2230-8210.183480.

5. Mohamed J, Nazratun Nafizah AH, Zariyantey AH, Budin SB. Mechanisms of diabetes-induced liver damage: The role of oxidative stress and inflammation. Sultan Qaboos Univ Med J. 2016;16(2):e132–e141. doi:10.18295/squmj.2016.16.02.002.

6. Sertorio MN, Souza ACF, Bastos DSS, et al. Arsenic exposure intensifies glycogen nephrosis in diabetic rats. Environ Sci Pollut Res. 2019;26(12):12459–12469. doi:10.1007/s11356-019-04597-1.

7. Levelt E, Gulsin G, Neubauer S, McCann GP. Diabetic cardiomyopathy: pathophysiology and potential metabolic interventions state of the art review. Eur J Endocrinol. 2018;178(4):R127–R139. doi:10.1530/EJE-17-0724.

8. Ullah A, Khan A, Khan I. Diabetes mellitus and oxidative stress—A concise review. Saudi Pharm J. 2016;24(5):547–553. doi:10.1016/j.jsps.2015.03.013.

9. Sociedade Brasileira de Diabetes. Diretrizes - Sociedade Brasileira de Diabetes 2017-2018.; 2017.

10. Silva E, Natali AÔJ, Silva MF, et al. Ventricular remodeling in growing rats with experimental diabetes: The impact of swimming training. Pathol Res Pract. 2013;209(10):618–626. doi:10.1016/j.prp.2013.06.009.

11. da Silva MF, Natali AJ, da Silva E, et al. Attenuation of Ca 2+ homeostasis, oxidative stress, and mitochondrial dysfunctions in diabetic rat heart: insulin therapy or aerobic exercise? J Appl Physiol. 2015;119(2):148–156. doi:10.1152/japplphysiol.00915.2014.

12. da Silva E, Natali AJ, da Silva MF, et al. Swimming training attenuates the morphological reorganization of the myocardium and local inflammation in the left ventricle of growing rats with untreated experimental diabetes. Pathol - Res Pract. 2016;212(4):325–334. doi:10.1016/j.prp.2016.02.005.

13. Babu PVA, Sabitha KE, Srinivasan P, Shyamaladevi CS. Green tea attenuates diabetes induced Maillard-type fluorescence and collagen cross-linking in the heart of streptozotocin diabetic rats. Pharmacol Res. 2007;55(5):433–440. doi:10.1016/j.phrs.2007.01.019.

14. Samarghandian S, Azimi-Nezhad M, Farkhondeh T. Catechin Treatment Ameliorates Diabetes and Its Complications in Streptozotocin-Induced Diabetic Rats. Dose-Response. 2017;15(1):155932581769115. doi:10.1177/1559325817691158.

15. Othman AI, El-Sawi MR, El-Missiry MA, Abukhalil MH. Epigallocatechin-3-gallate protects against diabetic cardiomyopathy through modulating the cardiometabolic risk factors, oxidative stress, inflammation, cell death and fibrosis in streptozotocin-nicotinamide-induced diabetic rats. Biomed Pharmacother. 2017;94:362–373. doi:10.1016/j.biopha.2017.07.129.

16. Latief U, Ahmad R. Herbal remedies for liver fibrosis: A review on the mode of action of fifty herbs. J Tradit Complement Med. 2018;8(3):352–360. doi:10.1016/j.jtcme.2017.07.002.

17. Roghani M, Baluchnejadmojarad T. Hypoglycemic and hypolipidemic effect and antioxidant activity of chronic epigallocatechin-gallate in streptozotocin-diabetic rats. Pathophysiology. 2010;17(1):55–59. doi:10.1016/j.pathophys.2009.07.004.

18. Li T, Liu J, Zhang X, Ji G. Antidiabetic activity of lipophilic (−)-epigallocatechin-3-gallate derivative under its role of α-glucosidase inhibition. Biomed Pharmacother. 2007;61(1):91–96. doi:10.1016/j.biopha.2006.11.002.

19. Anderson RA, Polansky MM. Tea Enhances Insulin Activity. J Agric Food Chem. 2002;50(24):7182–7186. doi:10.1021/jf020514c.

20. Zeng X, Tan X. Epigallocatechin-3-gallate and zinc provide anti-apoptotic protection against hypoxia/reoxygenation injury in H9c2 rat cardiac myoblast cells. Mol Med Rep. 2015;12(2):1850–1856. doi:10.3892/mmr.2015.3603.

21. Baluchnejadmojarad T, Roghani M. Chronic Oral Epigallocatechin-gallate Alleviates Streptozotocin-induced Diabetic Neuropathic Hyperalgesia in Rat: Involvement of Oxidative Stress. Iran J Pharm Res IJPR. 2012;11(4):1243–1253. http://www.ncbi.nlm.nih.gov/pubmed/24250559.

22. Waltner-Law ME, Wang XL, Law BK, Hall RK, Nawano M, Granner DK. Epigallocatechin Gallate, a Constituent of Green Tea, Represses Hepatic Glucose Production. J Biol Chem. 2002;277(38):34933–34940. doi:10.1074/jbc.M204672200.

23. Collins QF, Liu H-Y, Pi J, Liu Z, Quon MJ, Cao W. Epigallocatechin-3-gallate (EGCG), A Green Tea Polyphenol, Suppresses Hepatic Gluconeogenesis through 5′-AMP-activated Protein Kinase. J Biol Chem. 2007;282(41):30143–30149. doi:10.1074/jbc.M702390200.

24. Li Y, Zhao S, Zhang W, et al. Epigallocatechin-3-O-gallate (EGCG) attenuates FFAs-induced peripheral insulin resistance through AMPK pathway and insulin signaling pathway in vivo. Diabetes Res Clin Pract. 2011;93(2):205–214. doi:10.1016/j.diabres.2011.03.036.

25. Tang W, Li S, Liu Y, Huang M-T, Ho C-T. Anti-diabetic activity of chemically profiled green tea and black tea extracts in a type 2 diabetes mice model via different mechanisms. J Funct Foods. 2013;5(4):1784–1793. doi:10.1016/j.jff.2013.08.007.

26. Choo JJ. Green tea reduces body fat accretion caused by high-fat diet in rats through ␤-adrenoceptor activation of thermogenesis in brown adipose tissue. J Nutr Biochem. 2003;11:671–676. doi:10.1016/j.nutbio.2003.08.005.

27. Sae-tan S, Rogers CJ, Lambert JD. Voluntary exercise and green tea enhance the expression of genes related to energy utilization and attenuate metabolic syndrome in high fat fed mice. Mol Nutr Food Res. 2014;58(5):1156–1159. doi:10.1002/mnfr.201300621.

28. Stohs SJ, Badmaev V. A Review of Natural Stimulant and Non-stimulant Thermogenic Agents. Phyther Res. 2016;30(5):732–740. doi:10.1002/ptr.5583.

29. Türközü D, Tek NA. A minireview of effects of green tea on energy expenditure. Crit Rev Food Sci Nutr. 2017;57(2):254–258. doi:10.1080/10408398.2014.986672.

30. Yoneshiro T, Matsushita M, Hibi M, et al. Tea catechin and caffeine activate brown adipose tissue and increase cold-induced thermogenic capacity in humans. Am J Clin Nutr. 2017;105(4):873–881. doi:10.3945/ajcn.116.144972.

31. Islam MS, Choi H. Green tea, anti-diabetic or diabetogenic: A dose response study. BioFactors. 2007;29(1):45–53. doi:10.1002/biof.5520290105.

32. Rasheed NOA, Ahmed LA, Abdallah DM, El-Sayeh BM. Nephro-toxic effects of intraperitoneally injected EGCG in diabetic mice: involvement of oxidative stress, inflammation and apoptosis. Sci Rep. 2017;7(1):40617. doi:10.1038/srep40617.

33. Rasheed NOA, Ahmed LA, Abdallah DM, El-Sayeh BM. Paradoxical cardiotoxicity of intraperitoneally-injected epigallocatechin gallate preparation in diabetic mice. Sci Rep. 2018;8(1):7880. doi:10.1038/s41598-018-25901-y.

34. Silva MJ, Brodt MD, Lynch MA, et al. Type 1 Diabetes in Young Rats Leads to Progressive Trabecular Bone Loss, Cessation of Cortical Bone Growth, and Diminished Whole Bone Strength and Fatigue Life. J Bone Miner Res. 2009;24(9):1618–1627. doi:10.1359/jbmr.090316.

35. Locatto ME, Abranzon H, Caferra D, Fernandez M del C, Alloatti R, Puche RC. Growth and development of bone mass in untreated alloxan diabetic rats. Effects of collagen glycosylation and parathyroid activity on bone turnover. Bone Miner. 1993;23(2):129–144. doi:10.1016/S0169-6009(08)80049-9.

36. Perva-Uzunalić A, Škerget M, Knez Ž, Weinreich B, Otto F, Grüner S. Extraction of active ingredients from green tea (Camellia sinensis): Extraction efficiency of major catechins and caffeine. Food Chem. 2006;96(4):597–605. doi:10.1016/j.foodchem.2005.03.015.

37. Singleton VL, Rossi JA, Jr J. Colorimetry of Total Phenolics With Phosphomolybdic-Phosphotungstic Acid Reagents. Am J Enol Vitic. 1965;16(3):144–158.

38. Kim-Park WK, Allam ES, Palasuk J, Kowolik M, Park KK, Windsor LJ. Green tea catechin inhibits the activity and neutrophil release of Matrix Metalloproteinase-9. J Tradit Complement Med. 2016;6(4):343–346. doi:10.1016/j.jtcme.2015.02.002.

39. Re R, Pellegrini N, Proteggente A, Pannala A, Yang M, Rice-Evans C. Antioxidant activity applying an improved ABTS radical cation decolorization assay. Free Radic Biol Med. 1999;26(9-10):1231–1237. doi:10.1016/S0891-5849(98)00315-3.

40. Benzie IFF, Strain JJ. The Ferric Reducing Ability of Plasma (FRAP) as a Measure of “Antioxidant Power”: The FRAP Assay. Anal Biochem. 1996;239(1):70–76. doi:10.1006/abio.1996.0292.

41. Haidara MA, Mikhailidis DP, Rateb MA, et al. Evaluation of the effect of oxidative stress and vitamin E supplementation on renal function in rats with streptozotocin-induced Type 1 diabetes. J Diabetes Complications. 2009;23(2):130–136. doi:10.1016/j.jdiacomp.2008.02.011.

42. Sengupta P. The Laboratory Rat: Relating Its Age With Human’s. Int J Prev Med. 2013;4(6):624–630. http://www.ncbi.nlm.nih.gov/pubmed/23930179%0Ahttp://www.pubmedcentral.nih.gov/articlerender.fcgi?artid=PMC3733029.

43. Nery C da S, Pinheiro IL, Muniz G de S, de Vasconcelos DAA, de França SP, do Nascimento E. Murinometric evaluations and feed efficiency in rats from reduced litter during lactation and submitted or not to swimming exercise. Rev Bras Med do Esporte. 2011;17(1):49–55. doi:10.1590/S1517-86922011000100010.

44. Melo DS, Costa-Pereira L V., Santos CS, et al. Severe Calorie Restriction Reduces Cardiometabolic Risk Factors and Protects Rat Hearts from Ischemia/Reperfusion Injury. Front Physiol. 2016;7(APR):1–8. doi:10.3389/fphys.2016.00106.

45. Khan N, Mukhtar H. Tea polyphenols for health promotion. Life Sci. 2007;81(7):519–533. doi:10.1016/j.lfs.2007.06.011.

46. da Silva Pinto M. Tea: A new perspective on health benefits. Food Res Int. 2013;53(2):558–567. doi:10.1016/j.foodres.2013.01.038.

47. Sharangi AB. Medicinal and therapeutic potentialities of tea (Camellia sinensis L.) – A review. Food Res Int. 2009;42(5-6):529–535. doi:10.1016/j.foodres.2009.01.007.

48. Chung J-O, Yoo S-H, Lee Y-E, et al. Hypoglycemic potential of whole green tea: water-soluble green tea polysaccharides combined with green tea extract delays digestibility and intestinal glucose transport of rice starch. Food Funct. 2019;10(2):746–753. doi:10.1039/C8FO01936C.

49. Fu Q-Y, Li Q-S, Lin X-M, et al. Antidiabetic Effects of Tea. Molecules. 2017;22(5):849. doi:10.3390/molecules22050849.

50. Yan J, Zhao Y, Suo S, Liu Y, Zhao B. Green tea catechins ameliorate adipose insulin resistance by improving oxidative stress. Free Radic Biol Med. 2012;52(9):1648–1657. doi:10.1016/j.freeradbiomed.2012.01.033.

51. Wei K, Eckmanns T, Oppert M, et al. The Streptozotocin-Diabetic Chronic Complications Rat as a Model of the of Human Diabetes. Hear Lung Circ. 2003:1–20. doi:10.1067/mod.2000.104493.

52. Kobayashi Y, Suzuki M, Satsu H, et al. Green Tea Polyphenols Inhibit the Sodium-Dependent Glucose Transporter of Intestinal Epithelial Cells by a Competitive Mechanism. J Agric Food Chem. 2000;48(11):5618–5623. doi:10.1021/jf0006832.

53. Boyer S, Sharp PA, Debnam ES, Baldwin SA, Srai SKS. Streptozotocin diabetes and the expression of GLUT1 at the brush border and basolateral membranes of intestinal enterocytes. FEBS Lett. 1996;396(2-3):218–222. doi:10.1016/0014-5793(96)01102-7.

54. Wong TP, Debnam ES, Leung PS. Diabetes mellitus and expression of the enterocyte renin-angiotensin system: implications for control of glucose transport across the brush border membrane. Am J Physiol Physiol. 2009;297(3):C601–C610. doi:10.1152/ajpcell.00135.2009.

55. Kellett GL, Brot-Laroche E, Mace OJ, Leturque A. Sugar Absorption in the Intestine: The Role of GLUT2. Annu Rev Nutr. 2008;28(1):35–54. doi:10.1146/annurev.nutr.28.061807.155518.

56. Corpe CP, Basaleh MM, Affleck J, Gould G, Jess TJ, Kellett GL. The regulation of GLUT5 and GLUT2 activity in the adaptation of intestinal brush-border fructose transport in diabetes. Pflügers Arch - Eur J Physiol. 1996;432(2):192–201. doi:10.1007/s004240050124.

57. Al-Malki AL, El Rabey HA. The Antidiabetic Effect of Low Doses of Moringa oleifera Lam. Seeds on Streptozotocin Induced Diabetes and Diabetic Nephropathy in Male Rats. Biomed Res Int. 2015;2015:1–13. doi:10.1155/2015/381040.

58. Choi D, Piao Y, Yu S-J, et al. Antihyperglycemic and antioxidant activities of polysaccharide produced from Pleurotus ferulae in streptozotocin-induced diabetic rats. Korean J Chem Eng. 2016;33(6):1872–1882. doi:10.1007/s11814-016-0007-8.

59. Hwang H-J, Kim S-W, Lim J-M, et al. Hypoglycemic effect of crude exopolysaccharides produced by a medicinal mushroom Phellinus baumii in streptozotocin-induced diabetic rats. Life Sci. 2005;76(26):3069–3080. doi:10.1016/j.lfs.2004.12.019.

60. Herrero P, Peterson LR, McGill JB, et al. Increased Myocardial Fatty Acid Metabolism in Patients With Type 1 Diabetes Mellitus. J Am Coll Cardiol. 2006;47(3):598–604. doi:10.1016/j.jacc.2005.09.030.

61. Quesada I, Tudurí E, Ripoll C, Nadal Á. Physiology of the pancreatic α-cell and glucagon secretion: role in glucose homeostasis and diabetes. J Endocrinol. 2008;199(1):5–19. doi:10.1677/JOE-08-0290.

62. Slavin BG, Ong JM, Kern PA. Hormonal regulation of hormone-sensitive lipase activity and mRNA levels in isolated rat adipocytes. J Lipid Res. 1994;35(9):1535–1541. http://www.ncbi.nlm.nih.gov/pubmed/7806967.

63. Lu H. Enzymology of Methylation of Tea Catechins and Inhibition of Catechol-O-methyltransferase by (−)-Epigallocatechin Gallate. Drug Metab Dispos. 2003;31(5):572–579. doi:10.1124/dmd.31.5.572.

64. Shixian Q, VanCrey B, Shi J, Kakuda Y, Jiang Y. Green Tea Extract Thermogenesis-Induced Weight Loss by Epigallocatechin Gallate Inhibition of Catechol-O-Methyltransferase. J Med Food. 2006;9(4):451–458. doi:10.1089/jmf.2006.9.451.

65. Sorenson RL, Elde RP, Seybold V. Effect of Norepinephrine on Insulin, Glucagon, and Somatostatin Secretion in Isolated Perifused Rat Islets. Diabetes. 1979;28(10):899–904. doi:10.2337/diab.28.10.899.

66. Burcelin R, Eddouks M, Maury J, Kande J, Assan R, Girard J. Excessive glucose production, rather than insulin resistance, accounts for hyperglycaemia in recent-onset streptozotocin-diabetic rats. Diabetologia. 1995;38(3):283–290. doi:10.1007/BF00400632.

67. Vergès B. Lipid disorders in type 1 diabetes. Diabetes Metab. 2009;35(5):353–360. doi:10.1016/j.diabet.2009.04.004.

68. Livesey G, Elia M. Estimation of energy expenditure, net carbohydrate utilization, and net fat oxidation and synthesis by indirect calorimetry: evaluation of errors with special reference to the detailed composition of fuels. Am J Clin Nutr. 1988;47(4):608–628. doi:10.1093/ajcn/47.4.608.

69. Peronnet F, Massicotte D. Table of nonprotein respiratory quotient: an update. Can J Sport Sci. 1991;16:23–29.

70. Pujia A, Mazza E, Ferro Y, et al. Lipid Oxidation Assessed by Indirect Calorimetry Predicts Metabolic Syndrome and Type 2 Diabetes. Front Endocrinol (Lausanne). 2019;9(January):1–7. doi:10.3389/fendo.2018.00806.

71. Sae-tan S, Grove KA, Kennett MJ, Lambert JD. (−)-Epigallocatechin-3-gallate increases the expression of genes related to fat oxidation in the skeletal muscle of high fat-fed mice. Food Funct. 2011;2(2):111. doi:10.1039/c0fo00155d.

72. Sae-tan S, Rogers CJ, Lambert JD. Decaffeinated green tea and voluntary exercise induce gene changes related to beige adipocyte formation in high fat-fed obese mice. J Funct Foods. 2015;14:210–214. doi:10.1016/j.jff.2015.01.036.

73. Barja de Quiroga G. Brown fat thermogenesis and exercise: two examples of physiological oxidative stress? Free Radic Biol Med. 1992;13(4):325–340. http://www.ncbi.nlm.nih.gov/pubmed/1398216.

